# Characterization of a β-lactamase that contributes to intrinsic β-lactam resistance in *Clostridioides difficile*

**DOI:** 10.1101/630020

**Authors:** Brindar K. Sandhu, Adrianne N. Edwards, Sarah E. Anderson, Emily C. Woods, Shonna M. McBride

## Abstract

*Clostrididioides difficile* causes severe antibiotic-associated diarrhea and colitis. *C. difficile* is an anaerobic, Gram-positive spore former that is highly resistant to β-lactams, the most commonly prescribed antibiotics. The resistance of *C. difficile* to β-lactam antibiotics allows the pathogen to replicate and cause disease in antibiotic-treated patients. However, the mechanisms of β-lactam resistance in *C. difficile* are not fully understood. Our data reinforce prior evidence that *C. difficile* produces a β-lactamase, which is a common β-lactam resistance mechanism found in other bacterial species. We identified an operon encoding a lipoprotein of unknown function and a β-lactamase that was greatly induced in response to several classes of β-lactam antibiotics. An in-frame deletion of the operon abolished β-lactamase activity in *C. difficile* strain 630Δ*erm* and resulted in decreased resistance to the β-lactam ampicillin. We found that the activity of this β-lactamase, herein named BlaD, is dependent upon the redox state of the enzyme. In addition, we observed that transport of BlaD out of the cytosol and to the cell surface is facilitated by an N-terminal signal sequence. Our data demonstrate that a co-transcribed lipoprotein, BlaX, aids in BlaD activity. Further, we identified a conserved BlaRI regulatory system and demonstrated via insertional disruption that BlaRI controls transcription of the *blaXD* operon in response to β-lactams. These results provide support for the function of a β-lactamase in *C. difficile* antibiotic resistance, and reveal the unique roles of a co-regulated lipoprotein and reducing environment in β-lactamase activity.

**IMPORTANCE:** *Clostridioides difficile* is an anaerobic, gastrointestinal human pathogen. One of the highest risk factors for contracting *C. difficile* infection is antibiotic treatment, which causes microbiome dysbiosis. *C. difficile* is resistant to β-lactam antibiotics, the most commonly prescribed class of antibiotics. *C. difficile* produces a recently discovered β-lactamase, which cleaves and inactivates numerous β-lactams. In this study, we report the contribution of this anaerobic β-lactamase to ampicillin resistance in *C. difficile*, as well as the transcriptional regulation of the gene, *blaD*, by a BlaRI system. In addition, our data demonstrate co-transcription of *blaD* with *blaX*, which encodes a membrane protein of previously unknown function. Furthermore, we provide evidence that BlaX enhances β-lactamase activity in a portion of *C. difficile* strains. This study demonstrates a novel interaction between a β-lactamase and a membrane protein in a Gram-positive pathogen, and due to the anaerobic nature of the β-lactamase activity, suggests that more β-lactamases are yet to be identified in other anaerobes.

## INTRODUCTION

*Clostridioides difficile*, or *C. difficile*, is an anaerobic, Gram-positive, spore-forming bacterial pathogen that causes antibiotic-associated diarrhea (1-3). *C. difficile* infection, or CDI, can be severe, resulting in psuedomembranous colitis, intestinal rupture, and death. The Center for Disease Control (CDC) estimates that almost half a million people in the U.S. suffer from CDI per year, resulting in approximately 29,000 deaths per year (4). As a result, CDI cases add approximately $4.8 billion per year to U.S. healthcare costs (5). *C. difficile* was first linked to antibiotic-associated diarrhea in 1978, and antibiotic treatment is still one of the highest risk factors for CDI (2, 3). Antibiotic treatment results in gastrointestinal dysbiosis, eliminating important indigenous anaerobes, thereby allowing for *C. difficile* population expansion (6, 7). Antibiotic treatment of CDI is limited to the use of vancomycin, fidaxomicin, or metronidazole, due to the high resistance *C. difficile* exhibits for a wide array of antibiotics (8-10).

The most commonly prescribed class of antibiotics are the β-lactams, which comprise 62% of all prescribed antibiotics in the United States and are strongly associated with *C. difficile* infections (11-13). β-lactams are inhibitors of bacterial cell wall synthesis and are characterized by a four-membered core lactam ring (14). β-lactams are further classified into four groups based on adjoining structures: the penicillins, cephalosporins, monobactams, and carbapenems (15). All β-lactam antibiotics bind to, and thus disable, cell-wall synthesizers called penicillin-binding proteins (PBPs) of bacteria (16, 17). Since the introduction of β-lactams into modern medicine, multiple mechanisms of resistance to these antibiotics have been discovered in a variety of bacterial species. β-lactam resistance mechanisms include the production of β-lactamases, which hydrolyze the β-lactam ring and render the antibiotic ineffective, mutations acquired in PBPs that prevent binding of the β-lactams, reduced outer membrane permeability due to reduced porin expression, and efflux pumps, which prevent the antibiotic from reaching the cell wall (18-23).

The most common mechanism of β-lactam resistance occurs through the production of β-lactamase enzymes. Most of the characterized β-lactamases have been identified in Gram-negative species; in these bacteria, the β-lactamase is generally secreted into the periplasm, where the enzyme is concentrated, allowing for high levels of β-lactam resistance (24). Less common are the outer membrane-anchored β-lactamases, which may be further packaged into outer membrane vesicles, enabling the inactivation of nearby β-lactams (25-27). β-lactam resistance in Gram-positive bacteria, however, is more commonly conferred by the modification of the intended targets of the β-lactam, the penicillin-binding proteins (28). Still, β-lactamases do exist in Gram-positive bacteria (29-33). Although Gram-positive bacteria lack a periplasmic space, some species do produce membrane-bound β-lactamases (29, 34-37). A few of these enzymes are proteolytically cleaved, producing an exoenzyme that can be released from the membrane (31, 36, 38).

β-lactamase enzymes are classified into four classes: A, B, C, and D. Classes A, C, and D are serine hydrolases, while class B β-lactamases are metallohydrolases (18). Whereas β-lactamases of all classes have been discovered in Gram-negative bacteria, most Gram-positive β-lactamases belong to classes A or B (32). Class D β-lactamases were recently identified in Gram-positive bacteria, including one that is highly conserved among *C. difficile* isolates (33, 39). A recent study demonstrated that a β-lactamase in *C. difficile* confers resistance to the penicillin, cephalosporin, and monobactam class of β-lactams (39). According to the substrate profile of this enzyme, this β-lactamase belongs to the 2de functional group of β-lactamases (39, 40). The purpose of our study was to characterize the genetic organization, resistance contributions, biochemical activity, and regulation of the *C. difficile* β-lactamase. To accomplish this, we deleted the genes encoding the β-lactamase and the upstream predicted membrane protein in *C. difficile*, and examined the resulting resistance profiles and biochemical activity. Notably, we observed that the *C. difficile* β-lactamases are inactivated by oxygen, which has not been described for other class D β-lactamases. We also examined how this β-lactamase enzyme is transported, and detail its mechanism of regulation. We demonstrate that unlike other described β-lactamases, the *C. difficile* β-lactamase is co-transcribed with a membrane protein that facilitates β-lactamase processing and function. These results further our understanding of β-lactam resistance in *C. difficile*, which may expose approaches to prevent or treat β-lactam-associated CDI.

## MATERIALS AND METHODS

### Bacterial strains and growth conditions

Bacterial strains and plasmids used in this study are listed in **Table 1**. *Escherichia coli* was grown at 37°C in LB medium with 100 μg/mL ampicillin (Sigma-Aldrich) and 20 μg/mL chloramphenicol (Sigma-Aldrich) when necessary (41). *C. difficile* was grown anaerobically at 37°C as previously described (42) in brain heart infusion medium supplemented with 2% yeast extract (BHIS; Βecton Dickinson Company) or Mueller Hinton Broth (MHB; Difco) with 2 μg/mL thiamphenicol (Sigma-Aldrich), 3.125 – 60 μg/mL cefoperazone (Sigma-Aldrich), 0.25 – 2 μg/mL ampicillin, 0.125 – 1.5 μg/mL imipenem (US Pharmacopeia), 0.75 μg/mL vancomycin (Sigma-Aldrich), 75 μg/mL polymyxin B (Sigma-Aldrich), 1 mg/mL lysozyme (Fisher Scientific), 7.5 μg/mL nisin (MP Biomedicals), 2 μg/mL LL-37 (Anaspec), or 250 μg/mL kanamycin (Sigma-Aldrich) when specified.

**Table 1.**
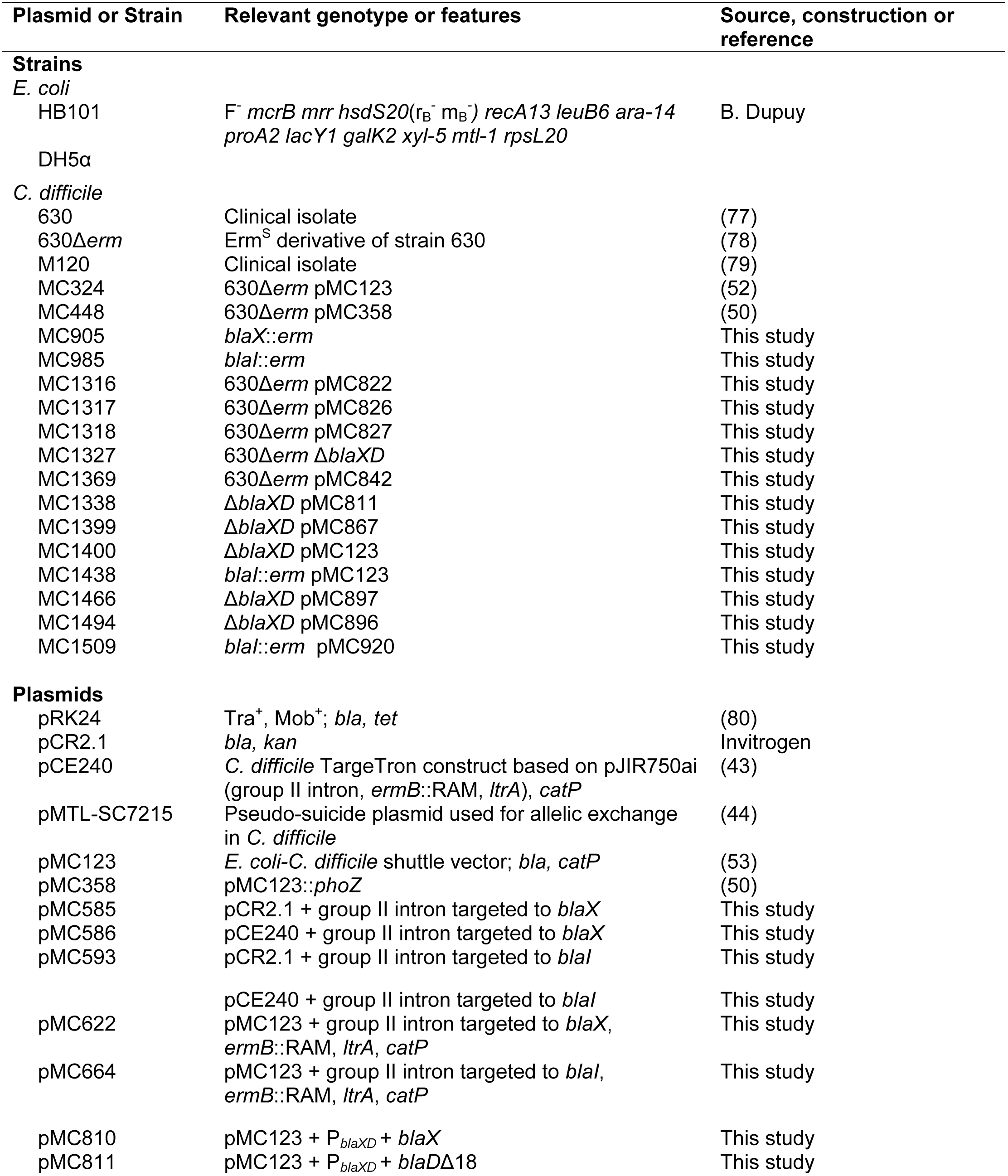
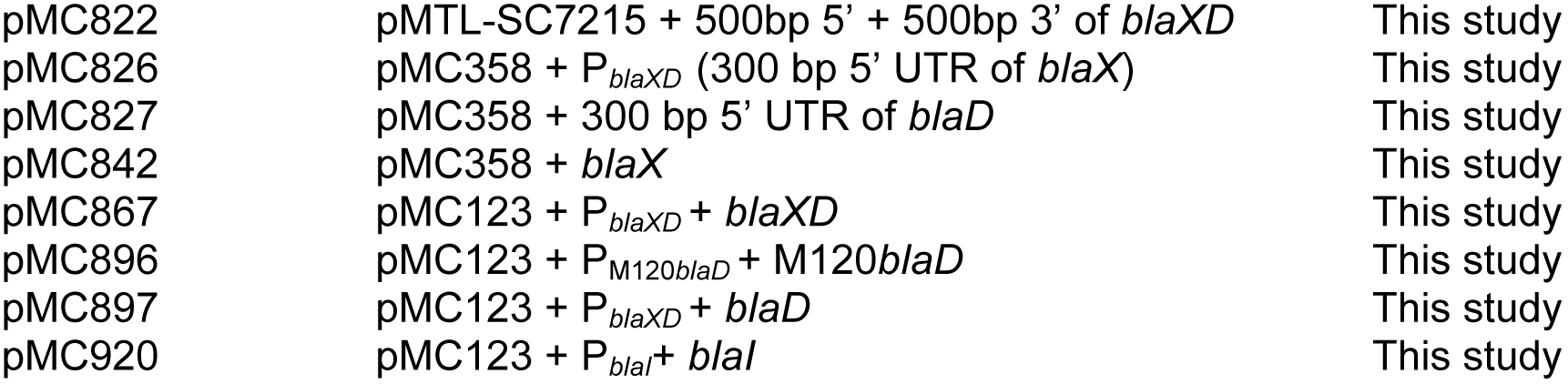
Bacterial Strains and plasmids.

### Strain and plasmid construction

The oligonucleotide primers used in this study are listed in **Table 2**. Primer design and the template for PCR reactions were based on *C. difficile* strain 630 (GenBank accession NC_009089.1), except for pMC896, which was based on strain M120 (GenBank accession FN665653.1).

**Table 2.**
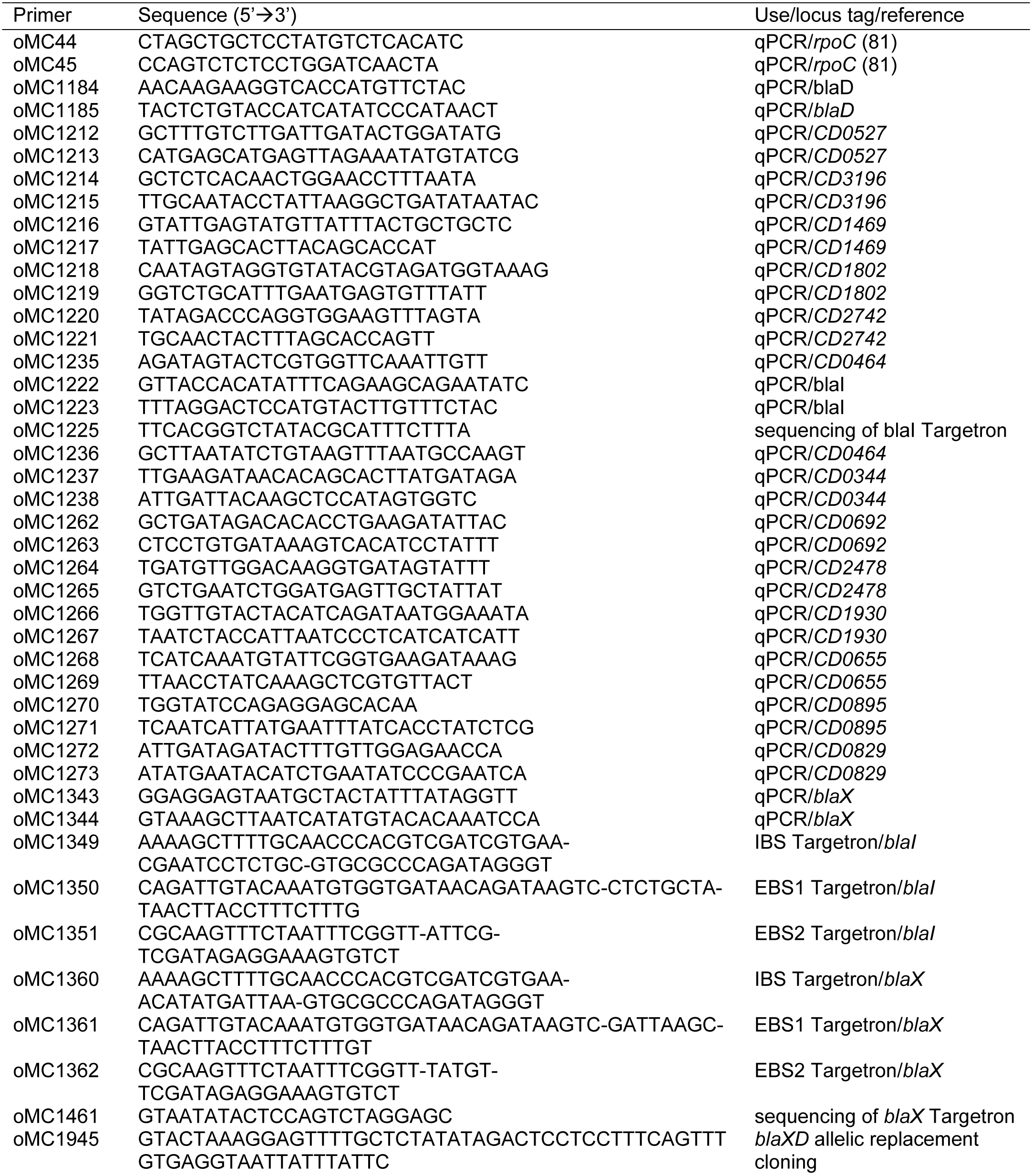
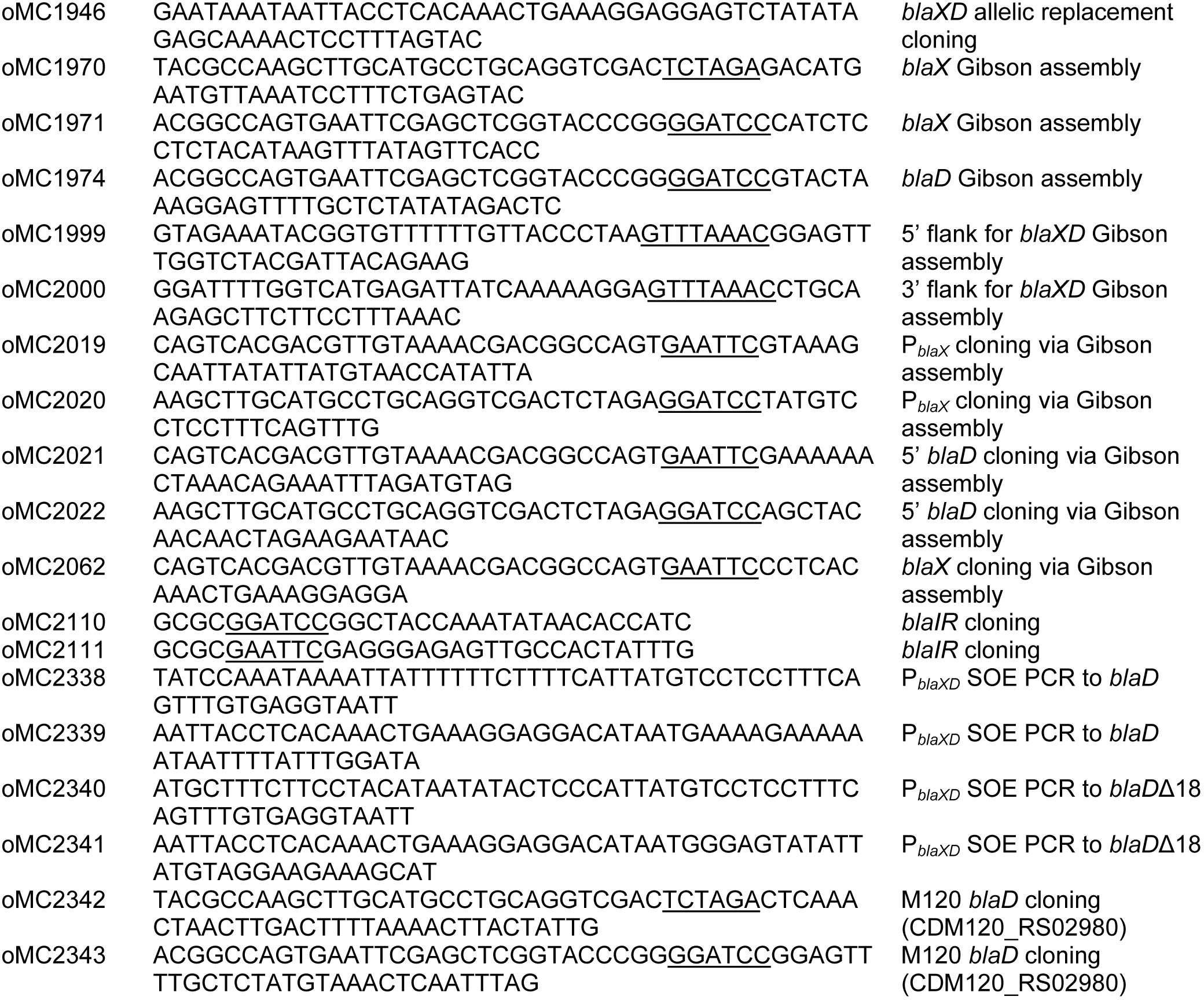
Oligonucleotides. Underlined nucleotides denote the restriction sites used for vector construction.

The *blaX*::*erm* and *blaI*::*erm* mutant strains were created by retargeting the Group II intron from pCE240 with the primers listed in **Table 2**, as previously described (43). To generate insertional disruptions, transconjugants were selected on 5 μg/mL erythromycin (Sigma-Aldrich), and 50 μg/mL kanamycin (Sigma-Aldrich) to select against *E. coli*.

The Δ*blaXD* mutant strain was created using a pseudo-suicide plasmid technique, as described previously, with slight variation (44). Briefly, 500 bp regions homologous to the 5’ and 3’ ends of the *bla* operon were amplified and Gibson assembled into the *Pme*I site of plasmid pMTLSC7215 to create plasmid pMC822. The plasmid was purified using a miniprep kit (Zymo Research), transformed into *E. coli* strain HB101 pRK24, and introduced into *C. difficile* by conjugation. *C. difficile* harboring the plasmid were selected on BHIS agar containing 15 μg/mL thiamphenicol, streaked onto BHIS agar, and subsequently on BHIS agar with 15 μg/mL thiamphenicol and 100 μg/mL kanamycin to force plasmid integration and counterselect against *E. coli*. A clone that screened positive for two crossover events was streaked to purity on BHIS agar for three more passages and the loss of plasmid was confirmed via sensitivity to 5 μg/mL thiamphenicol on BHIS agar.

Detailed construction of plasmids can be found in **Figure S1**. Plasmids were transferred to *C. difficile* as previously described, with slight variation (45, 46). Briefly, plasmids were chemically transformed into *E. coli* strain HB101 pRK24 and mated with *C. difficile* on agar plates for 48 h. Transconjugants were selected on BHIS agar containing 10 μg/mL thiamphenicol for plasmid selection and 100 μg/mL kanamycin to counterselect against *E. coli*.

### Nitrocefin hydrolysis disk assays

β-lactamase activity was assessed by hydrolysis of nitrocefin, a chromogenic cephalosporin (Sigma-Aldrich). Briefly, *C. difficile* was grown overnight in BHIS to log phase, then diluted to an OD_600_ of 0.05 in BHIS medium with or without 2 μg/mL ampicillin. Cultures were grown to an OD_600_ of 0.45-0.55, and 1 mL of culture was collected and centrifuged for 3 minutes at 21,130 rcf. All but approximately 30 μL of the supernatant was decanted, the pellets were resuspended, and the cells were spotted onto a nitrocefin disk. The disks were incubated aerobically or anaerobically for 2 h at 37°C, as noted.

### Nitrocefin liquid hydrolysis assays

β-lactamase activity was determined for complemented strains via anaerobic liquid nitrocefin assays, as previously reported, with some modifications (47). Briefly, *C. difficile* was grown overnight in BHIS with 2 μg/mL thiamphenicol to log phase, then diluted to an OD_600_ of 0.05 in BHIS medium with 2 μg/mL thiamphenicol and 2 μg/mL ampicillin. Cultures were grown to an OD_600_ of 0.45 – 0.55, 1 mL of culture was collected (in duplicate), and cells centrifuged for 3 min at 21,130 rcf. For whole cell reactions, supernatant was transferred to a fresh tube, and nitrocefin (BioVision) was added to supernatant or whole cell suspensions at a final concentration of 50 μM. For lysed cell reactions, pelleted cells were frozen at −20°C until use. Pellets were resuspended in lysis buffer (100 mM sodium phosphate + 50 mM sodium bicarbonate, pH 7.4), and DTT (Fisher Scientific) was added to each sample for a final concentration of 0.2 mM. Lysed samples were subjected to six freeze-thaw cycles (2 min in dry ice/Ethanol bath, 3 min at 37°C). 0.2 mL of the lysate was transferred to a fresh tube (designated ‘lysate’). The remaining volumes of samples were pelleted by centrifugation for 30 min at 21,130 rcf at 4°C, and then filtered via 0.22 μM syringe filters (BD Biosciences). 0.2 mL of this solution (designated ‘lysate filtrate’) was transferred to a fresh tube. Equal volumes of lysis buffer were added to each sample. Nitrocefin was added at a final concentration of 50 μM to bring the sample volume to 1 mL and samples were incubated anaerobically at 37°C for up to 7 minutes. Reactions were quenched by adding 100 μL of 1 M NaCl and immediately placed on ice. Samples were centrifuged for 3 min at 21,130 rcf to clear cell debris. The entire assay was performed anaerobically until this point. 300 μL of each supernatant was applied to a 96-well flat-bottom plate, and the OD_490_ was recorded with a BioTek microplate reader. β-lactamase units were calculated by the following equation: (OD_490_ * 1000) / (OD_600_ * time in min * vol of cells in mL), where OD_600_ is the value at the time of collection and the time is the number of minutes between the addition of nitrocefin and adding 1 M NaCl. Lysate results were normalized to the amount of lysate supernatant used. Time course experiments were run to confirm the linearity of the reaction. Results reported are the mean of at least three independent experiments.

### Minimal Inhibitory Concentration determination (MIC)

β-lactam susceptibility of *C. difficile* was determined as described previously (48). Briefly, active *C. difficile* cultures were diluted in Mueller Hinton Broth (MHB; BD Difco) to an OD_600_ of 0.1, which were grown to an OD_600_ of 0.45, and further diluted 1:10 in MHB. 15 μL of this diluted culture (∼5×10^5^ CFU/mL) was plated in a pre-reduced 96-well round bottom polystyrene plate that contained 135 μL of MHB with appropriate β-lactams in each well. The MIC was determined as the concentration at which there was no visible growth after 24 hours of anaerobic incubation at 37°C.

### Alkaline phosphatase activity assays

Alkaline phosphatase activity assays in *C. difficile* were performed as described previously, with minor modifications to the original published assay (49, 50). Briefly, *C. difficile* cultures were grown anaerobically at 37 °C overnight in BHIS with thiamphenicol (2 μg/mL) to log phase, then diluted to an OD_600_ of 0.05 in 10 mL BHIS with thiamphenicol. 1 mL of cells was collected in duplicate when the OD_600_ reached 0.5. Cells were centrifuged at 21,130 rcf for 3 min and the pellets were stored in −20°C at least overnight. For the assay, cell pellets were thawed and resuspended in 500 μL of cold wash buffer (10 mM Tris pH 8.0 + 10 mM MgSO_4_) and pelleted for 3 min at 21,130 rcf. Alkaline phosphatase assays were performed as previously described (50) without the addition of chloroform (51). The OD_550_ (cell debris) and OD_420_ (pNP cleavage) were measured in a BioTek microplate reader. Values were averaged between the triplicate wells, and then between duplicate technical samples. AP units were calculated as ((OD_420_ – (1.75* OD_550_)) * 1000) / (OD_600_ * time), where OD_600_ is the value at the time of collection. Results reported are the average between three independent experiments.

### Quantitative reverse transcription PCR analysis (qRT-PCR)

Actively growing *C. difficile* were diluted to an OD_600_ of 0.02 in 10 – 25 mL BHIS with appropriate antibiotic and grown to log phase. RNA was isolated as described previously (45, 52). Briefly, 3 mL samples were taken at an OD_600_ of 0.45 – 0.55, mixed with 3 mL ice-cold 1:1 acetone:ethanol, and stored immediately in −80°C. RNA was isolated (Qiagen RNeasy kit), treated for contaminating DNA (Invitrogen TURBO DNA-free kit), and RNA was reverse-transcribed into cDNA (Bioline Tetro cDNA synthesis kit). cDNA samples were used for qPCR (Bioline SensiFAST SYBR and Flourescein kit) in technical triplicates on a Roche Lightcycler 96 as described previously (53). Results are presented as the means and standard errors of the means for three biological replicates. Statistical significance was determined using a one-way ANOVA, followed by Dunnett’s multiple-comparison test (GraphPad Prism v6.0).

## RESULTS

### *C. difficile* produces an inducible, anaerobic β-lactamase

*C. difficile* was recently reported to produce a β-lactamase that can cleave β-lactam antibiotics (39). We further investigated the regulation and potential inducibility of *C. difficile* β-lactamase activity and examined the environmental conditions required for its function. Two diverse strains of *C. difficile*, 630Δ*erm* (ribotype 012) and R20291 (ribotype 027), were grown in the presence or absence of cefoxitin, a cephalosporin, and applied to a membrane disk impregnated with nitrocefin, a chromogenic cephalosporin. As shown in **Figure 1**, both strains of *C. difficile* grown in the presence of cefoxitin caused a color change from yellow to red, indicating cleavage of nitrocefin. In the absence of cefoxitin, neither strain demonstrated observable nitrocefin cleavage. These results suggested that *C. difficile* produces a β-lactamase that is inducible by β-lactams and is present in diverse strains. During optimization of these assays, we observed markedly higher β-lactamase activity under anaerobic conditions, suggesting that this activity was impaired by oxygen. Indeed, when the nitrocefin assay was performed in the presence of oxygen, the disk did not change color, indicating a loss of β-lactamase activity. These results demonstrate that *C. difficile* strains produce an inducible β-lactamase, and that the activity of this enzyme is quenched by oxygen.

**Figure 1.**
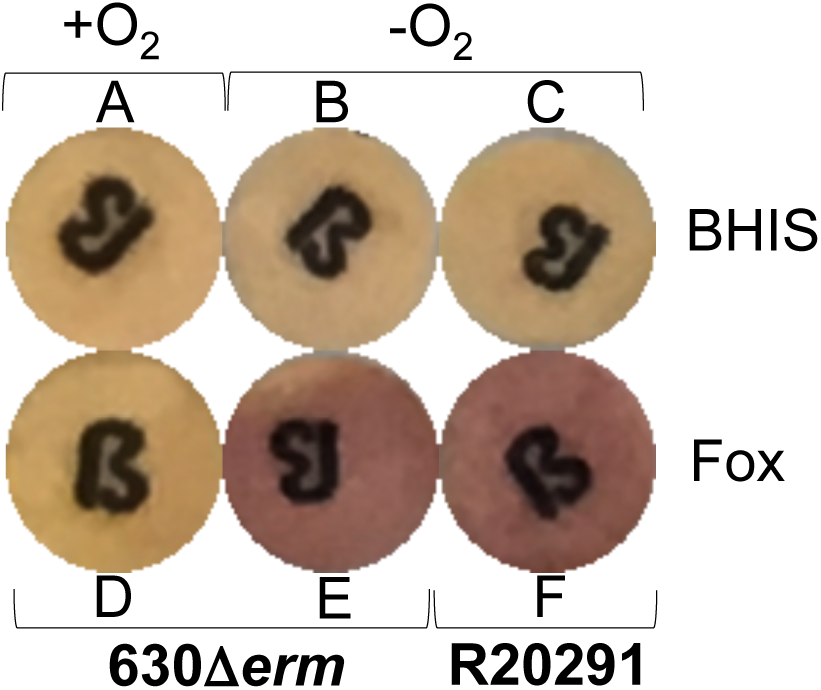
*C. difficile* strains exhibit inducible, anaerobic β-lactamase activity. Hydrolysis of the chromogenic cephalosporin nitrocefin was assessed for strains 630Δ*erm* (**A, B, D, E**) and R20291 (**C, F**). Strains were grown for ∼24 h on BHIS agar (**A-C**) or BHIS agar + 75 µg/ml cefoxitin (Fox; **D-F**). Cells were resuspended in water and incubated aerobically (**A, D**) or anaerobically (**B, C, E, F)** on nitrocefin disks. Color change from yellow to red indicates cleavage of nitrocefin.

### *CD0458* encodes the putative class D β-lactamase, BlaD

Based on the observed induction of β-lactamase activity, we hypothesized that the expression of one or more putative β-lactamases would be induced upon exposure to β-lactams. To test this, *C. difficile* strain 630Δ*erm* was grown in the presence of three classes of β-lactams: cefoperazone (a cephalosporin), ampicillin (a penicillin), and imipenem (a carbapenem). Using qRT-PCR, we measured the gene expression for 17 putative β-lactamases identified in the *C. difficile* genome (8, 54, 55). **Figure S2** demonstrates that the expression of only one of these genes, *CD0458*, was significantly induced upon exposure to each of the three types of β-lactams. This result supports a previously reported hypothesis, as *CD0458* was recently identified as a β-lactamase in *C. difficile* (39). This induction suggested that expression of *CD0458* confers the β-lactamase activity that we previously observed. The expression of the homologous gene in *C. difficile* strain R20291 was also greatly induced by these three β-lactams (CDR20291_0399, 99% identity; **Figure S2**). *CD0458* is analogous to two loci described recently by Toth *et al*. as *cdd1* and *cdd2* (39). However, based on the high similarity of the previously described Cdd1 and Cdd2 proteins, the existence of other genes already annotated as *cdd, cdd2*, and *cdd3* in strain 630 (56, 57), and the sequence similarity of the CD0458/CDR20291_0399 proteins to class D β-lactamases, we renamed the locus *blaD*.

### *CD0457* encodes a putative membrane protein, BlaX, which is co-transcribed with *blaD*

Analysis of the region surrounding *blaD* revealed the presence of another gene*, CD0457*, which appeared to be part of an operon with *blaD*. **Figure 2A** illustrates the putative *bla* operon, in which *CD0457* is located 27 nucleotides upstream of the start codon of *CD0458*. To determine if expression of *CD0457* is similarly induced upon β-lactam exposure, we measured transcription of *CD0457* in *C. difficile* strain 630Δ*erm* upon exposure to cefoperazone, ampicillin, and imipenem. **Figure 2B** demonstrates that expression of *CD0457* is comparably induced upon exposure to all three β-lactams. This co-regulation by β-lactams strongly suggested that *CD0457* is co-transcribed with *CD0458* and that the CD0457 predicted membrane protein product could play a role in the β-lactam resistance. The expression of the homologous gene in *C. difficile* strain R20291 was also comparably induced upon exposure to these β-lactams, indicating a similar organization in divergent strains (**Figure S3**).

**Figure 2.**
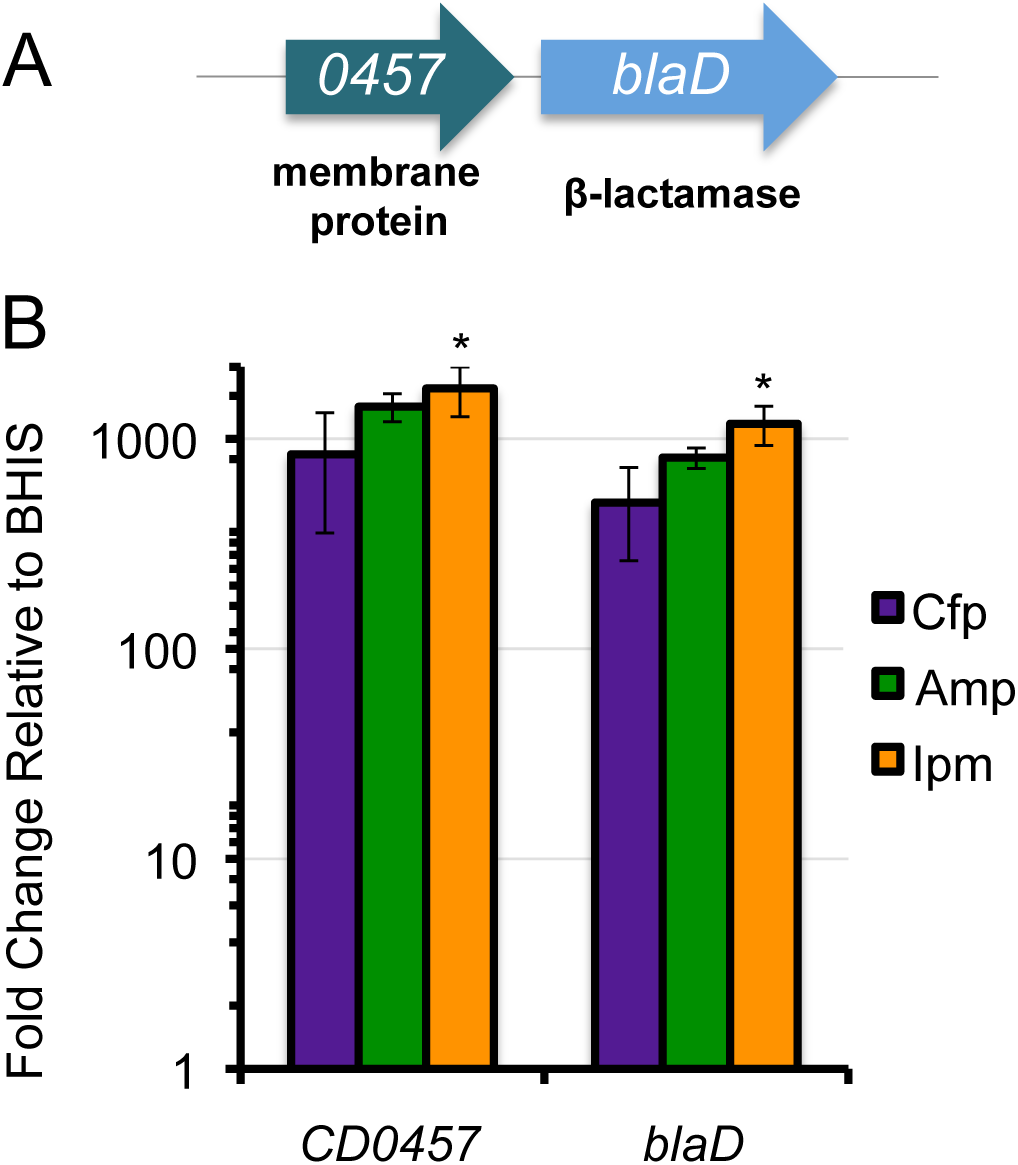
**The putative β-lactamase, *CD0458*, and the upstream gene, *CD0457* are induced by β-lactams. A**) The putative β-lactamase gene *CD0458* is 27 bp downstream of the predicted membrane protein, *CD0457.* **B**) Relative expression of each gene was measured via qRT-PCR. *C. difficile* strain 630Δ*erm* was grown to mid-log in BHIS medium supplemented with sub-inhibitory concentrations of β-lactams (Cfp: cefoperazone 50 µg/mL; Amp: ampicillin 2 µg/mL; Ipm: imipenem 1.5 µg/mL). mRNA levels are normalized to expression levels in BHIS alone. Columns represent the means +/− SEM from three independent replicates. Data were analyzed by one-way ANOVA with Dunnett’s multiple comparisons test, compared to no antibiotic. Adjusted *P* values indicated by *≤0.05.

To determine if the *CD0457* and *blaD* genes are part of a single cistronic unit, we assessed the linkage of these transcripts by amplifying the region between *CD0457* and *blaD* from cDNA generated after exposure of *C. difficile* strains 630Δ*erm* and R20291 to ampicillin (**Figure S4A**). **Figure S4B** illustrates the results of the PCR from cDNA that generated a product of 1 kb, which matches the genomic DNA product from the same strain. These data demonstrate that the transcription of *CD0457* and *blaD* are linked, indicating that they comprise a monocistronic unit. Since *CD0457* and *blaD* form an operon and the function of CD0457 is unknown, we named the *CD0457* gene *blaX*.

To further define the transcriptional organization of the *bla* operon, we examined promoter activity within the *bla* locus. Potential promoter activity was measured for putative promoter regions within the locus using *phoZ* reporter fusions, which produce alkaline phosphatase (50). As illustrated in **Figure 3**, regions of 300 nucleotides directly upstream of the start codons of *blaX* or *blaD* were fused to *phoZ* and expressed in *C. difficile*. The results of these reporter assays indicate that the region 300 nucleotides upstream of *blaX*, but not the region 300 nucleotides upstream of *blaD*, is able to promote transcription, resulting in measurable activity. To confirm the absence of a cryptic *blaD* promoter located within the *blaX* coding region, the entire region from the translational start of *blaX* to the start codon of *blaD* was also examined for possible promoter activity. However, no transcriptional activity was observed from this region (**Figure 3**). The only segment that produced significant and inducible activity contained the region upstream of the *blaX* coding sequence, strongly suggesting that soley this region drives *blaX* and *blaD* expression.

**Figure 3.**
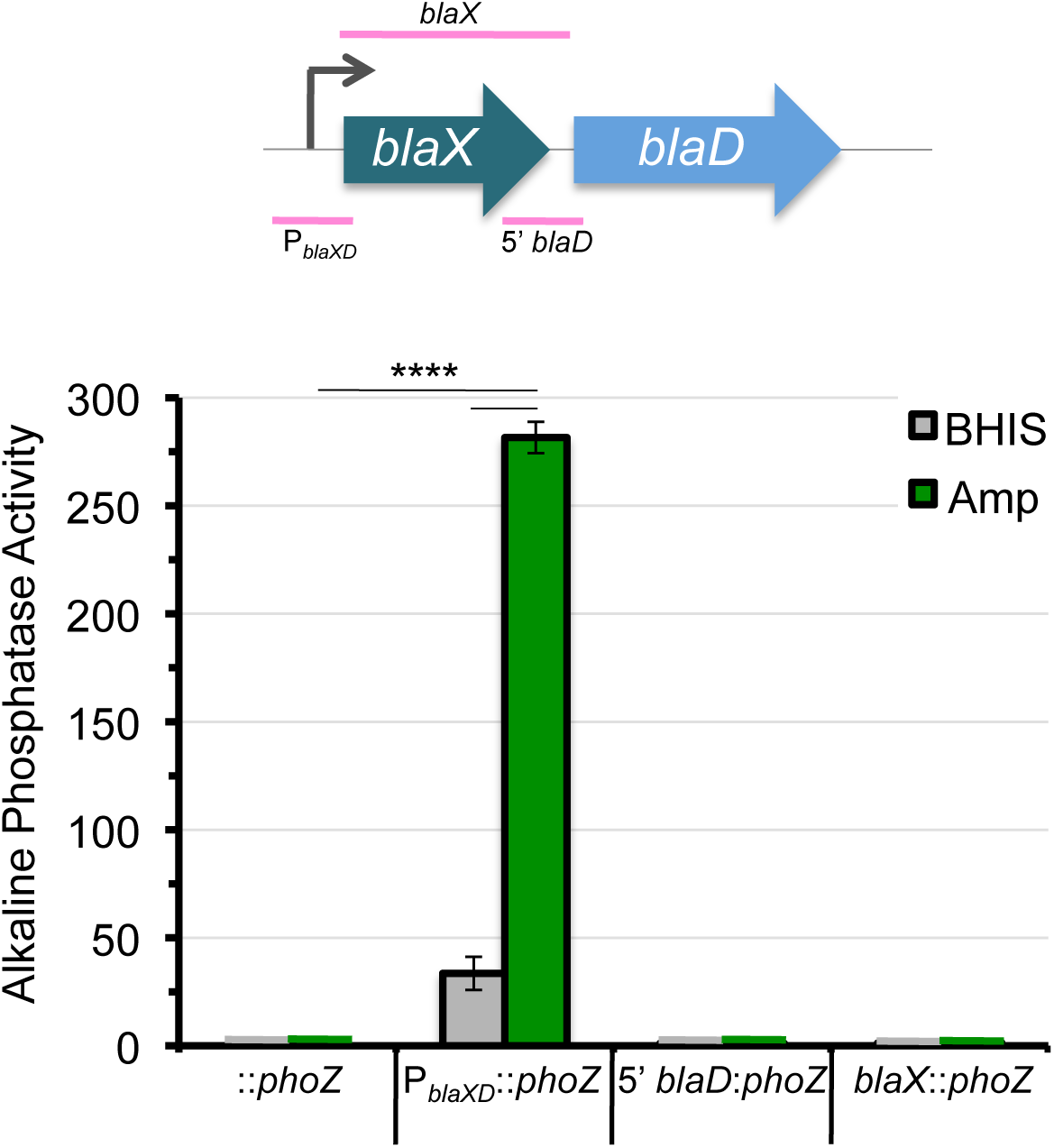
Alkaline phosphatase activity from P*_blaXD_::phoZ* is induced in the presence of ampicillin. *C. difficile* 630Δ*erm* cultures were grown to an OD_600_ of ∼0.5 in BHIS with 2 µg/mL thiamphenicol for plasmid maintenance in the presence or absence of 2 µg/mL ampicillin. Strains: MC448 (::*phoZ* - empty vector); MC1317 (P*_blaXD_*::*phoZ)*; MC1318 (5’ *blaD*::*phoZ)*; MC1369 (*blaX*::*phoZ)*. The means and standard errors of the means of three biological replicates are shown. Data were analyzed by one-way ANOVA with Dunnett’s multiple comparison test. Adjusted *P* value indicated by ****<0.0001.

### The *bla* operon contributes to ampicillin resistance in *C. difficile*

Notably, 36% of complete *C. difficile* genomes contain a homolog of *blaX*. Other sequenced genomes simply contain the same promoter and *blaD* region without the membrane protein. The membrane protein only shares approximately 23-40% amino acid identity to uncharacterized proteins found in a handful of other bacterial species. Thus, the function of this membrane protein cannot be inferred from other systems. To define the roles of BlaX and BlaD in β-lactam resistance and in β-lactamase activity, we created mutants of the 630Δ*erm* strain with an insertional mutation in the *blaX* gene (MC905) or complete deletion of the *blaX-blaD* locus (MC1327). Compared to the parent strain, *blaX*::*erm* displayed decreased, but still inducible *blaD* expression (**Figure S5**). Although *blaX* transcription is measurable in the *blaX*::*erm* mutant, the product is presumably non-functional because of the insertional mutation. We confirmed that neither the *blaX* nor the *blaD* transcript was expressed in the Δ*blaXD* mutant (**Figure S5**).

Based on the induction of β-lactamase activity and the induction of the *bla* operon by β-lactams, we hypothesized that deletion of the operon would reduce *C. difficile* resistance to β-lactams. As shown in **Figure 4**, we performed growth curves with the Δ*blaXD* and *blaX::erm* strains in cefoperazone, ampicillin, and imipenem to measure the contribution of the *bla* operon to β-lactam resistance in *C. difficile*. While the deletion of *blaX* and *blaD* did not significantly affect growth in cefoperazone, Δ*blaXD* and *blaX*::*erm* growth was impaired in ampicillin compared to the parent strain. These data suggest that the *bla* operon contributes to ampicillin resistance in *C. difficile*. Interestingly, the deletion of *blaX* and *blaD* improved growth in imipenem, supporting the finding by Toth *et al*. that BlaD binds to, but does not hydrolyze imipenem (39).

**Figure 4.**
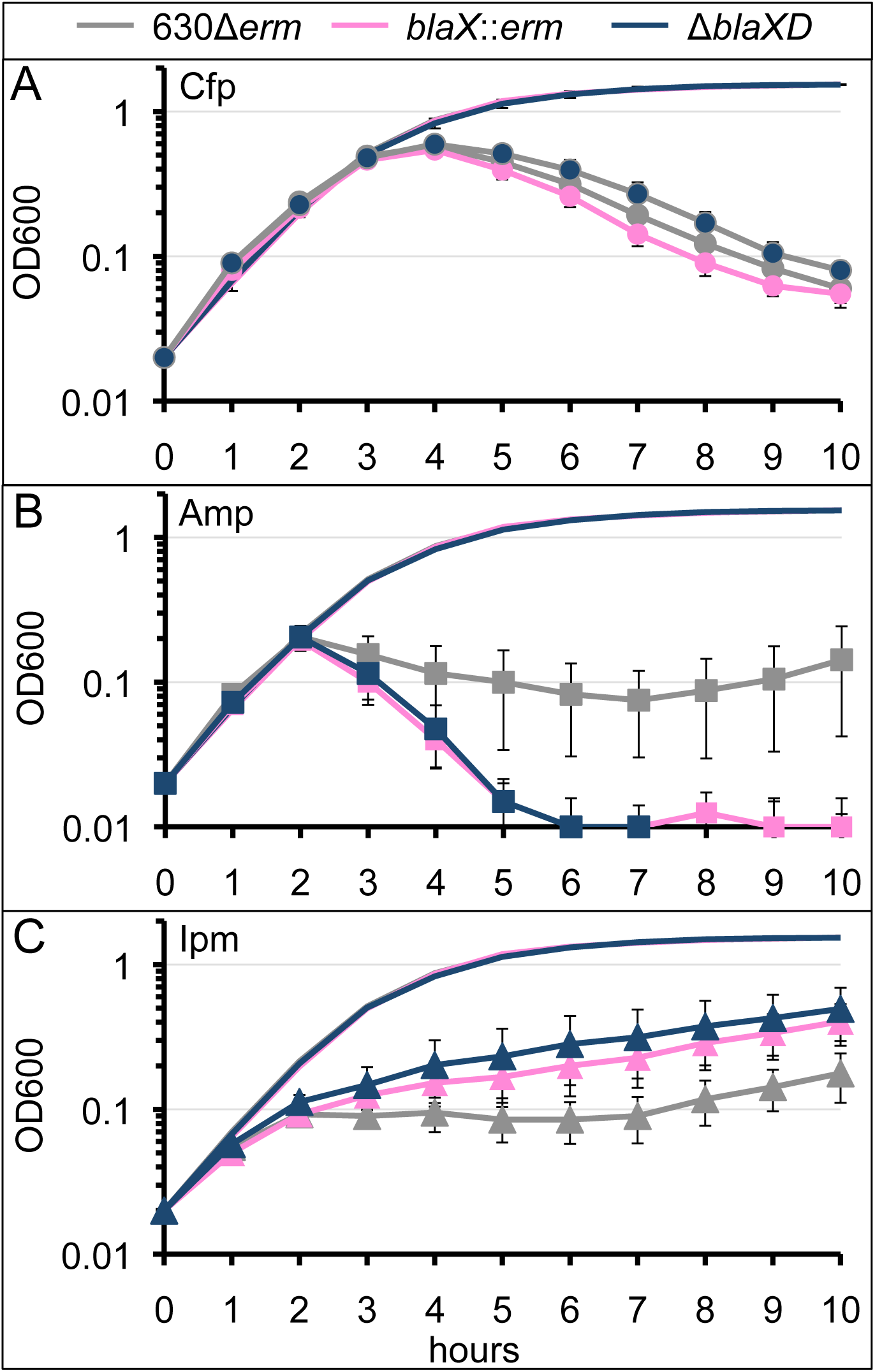
*blaX* and *blaD* contribute to β-lactam resistance in *C. difficile*. *C. difficile* strains 630Δ*erm* (green), *blaX*::*erm* (MC905; pink), and Δ*blaXD* (MC1327; blue) were grown to mid-log, backdiluted to OD 0.05, and grown in BHIS supplemented with **A**) Cfp: cefoperazone 60 μg/mL, **B**) Amp: ampicillin 4 μg/mL, or **C**) Ipm: imipenem 2 μg/mL. Lines represent the means +/− SEM from four independent replicates. Data were analyzed by one-tailed paired Student’s *t-*test, compared to 630Δ*erm*. No statistically significant differences found.

Antibiotic resistance in clinically relevant bacteria is often characterized by minimum inhibitory concentrations (MIC) of antibiotics. To further define the contribution of *blaX* and *blaD* to β-lactam resistance in *C. difficile*, we measured the MIC of β-lactams in 630Δ*erm*, Δ*blaXD*, and *blaX*::*erm*. Although the parent strain grew better in ampicillin, the MICs for both cefoperazone and ampicillin were similar in all three strains (**Table S1**), and higher for 630Δ*erm* in imipenem, indicating a modest difference in resistance values.

### The *bla* operon encodes the only functional β-lactamase in *C. difficile*

Although *blaD* was the only annotated β-lactamase induced by β-lactams (**Figure 1**), it was plausible that another β-lactamase existed in *C. difficile*. To determine if the *bla* operon encodes the only β-lactamase in *C. difficile*, we measured the β-lactamase activity of Δ*blaXD* in a nitrocefin hydrolysis assay. As shown in **Figure 5A**, no apparent β-lactamase activity was observed for the Δ*blaXD* strain. In comparison, the *blaX*::*erm* strain exhibits a slight change in color to a light pink, indicating that this mutant does not fully abolish production and activity of the β-lactamase, which is in agreement with the decrease in *blaD* gene expression observed for this strain (**Figure S5**). These results strongly suggest that *blaD* encodes the only functional β-lactamase in *C. difficile*.

**Figure 5.**
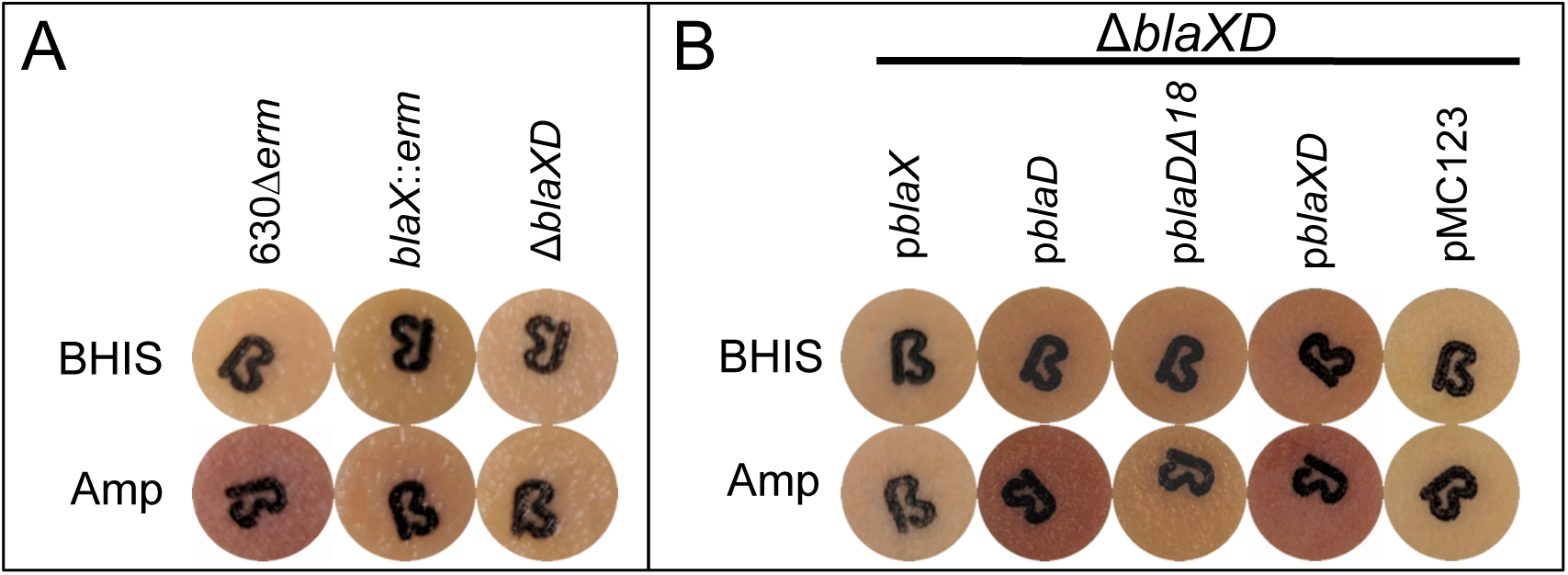
The N-terminus of BlaD is necessary for β-lactamase secretion, independent of BlaX. Hydrolysis of the chromogenic cephalosporin nitrocefin was assessed for **A**) strains 630Δ*erm*, *blaX*::*erm* (MC905), and Δ*blaXD* (MC1327) and **B**) strain Δ*blaXD* complemented with *blaX* and/or *blaD*, expressed from their native promoter. Strains were grown anaerobically to mid-log in BHIS medium (with 2 µg/mL thiamphenicol for plasmid maintenance in **B**) +/− 2 µg/mL ampicillin and pelleted. Cell pellets in ∼30 µL of remaining media were incubated anaerobically on nitrocefin disks for 2 h. Color change from yellow to red indicates cleavage of nitrocefin.

### The *bla* operon exhibits high level, dose-dependent expression in β-lactams

The induction of both *blaX* and *blaD* by β-lactams suggested that these genes are important for β-lactam resistance in *C. difficile*. To determine whether these genes could be induced by other cell wall targeting antimicrobials or if the induction is specific to β-lactam exposure, we measured the levels of gene expression for *C. difficile* strain 630Δ*erm* in various cell wall targeting antibiotics (vancomycin, polymyxin B, and lysozyme) and cationic antimicrobial peptides (nisin and LL-37), as well as a ribosome-targeting antibiotic (kanamycin). **Figure S6** shows that expression of *blaX* and *blaD* were induced in the presence of kanamycin and polymyxin B. However, these levels of expression are not statistically significant and were less than 3% of the levels seen for expression after β-lactam exposure, suggesting that the high levels of induction of *blaX* and *blaD* are specific to β-lactams.

Although the levels of *blaX* and *blaD* induction were high in all three β-lactams, expression varied greatly between each β-lactam. These results suggested that the level of induction of the *bla* operon is dependent upon the type of β-lactam *C. difficile* is exposed to and could be dose-dependent. To determine if the *bla* operon exhibits dose-dependent expression in β-lactams, we measured the relative expression of *blaX* and *blaD* in the 630Δ*erm* strain in varying concentrations of cefoperazone, ampicillin, and imipenem. **Figure S7** shows that the *bla* operon did indeed exhibit dose-dependent induction by β-lactams and that the response was different for the various classes of β-lactams. In increased concentrations of cefoperazone, induction of the *bla* operon trended downward, whereas expression trended upward in increased concentrations of ampicillin. Expression of the *bla* operon was high in all concentrations of imipenem, exhibiting only a modest increase in expression as the concentration of imipenem was increased. Furthermore, the level of induction of the *bla* operon was high even at concentrations of β-lactams far below the MIC (0.03125x MIC of cefoperazone, 0.125x of ampicillin, and 0.0625x MIC of imipenem). These results suggest that *bla* expression is controlled in a dose-dependent manner specific to the class of β-lactam administered.

### BlaX is not necessary for BlaD activity

Of the 72 genomes retrieved from a *blaD* BLASTn search of *C. difficile*, 42 strains encode the upstream putative membrane protein, suggesting that the membrane protein BlaX may be important for β-lactamase activity in some strains, but not in others. To determine the importance of the membrane protein, we first assessed β-lactamase activity in *C. difficile* strain M120, which lacks a homolog of *blaX*. The BlaD enzyme from strains M120 and 630Δ*erm* are highly similar, but the 4% variability clearly lies within the N-termini of these proteins (**Figure S8A**). As shown in **Figure S8B**, strain M120 does exhibit β-lactamase activity. The variability in the amino acid sequence of these two enzymes may be due to differences in signal sequence recognition, but a potential interaction with another protein cannot be ruled out.

As the function of BlaX was not immediately apparent, we examined whether BlaX is necessary to observe the β-lactamase activity of BlaD in strain 630Δ*erm*. To test this, we complemented the Δ*blaXD* strain with *blaX* and/or *blaD* in trans. The nitrocefin disk assays in **Figure 5B** demonstrate that expression of *blaD* alone can restore β-lactamase activity in the Δ*blaXD* mutant, indicating that BlaD can act independently of BlaX, despite the co-transcription of these two genes. This result is further supported by the observation that the *blaX*::*erm* strain exhibits some β-lactamase activity (**Figure 5A**).

### BlaD contains a predicted signal sequence and is associated with the cell membrane

A common characteristic of β-lactamases is an N-terminal signal sequence that directs the protein out of the cytoplasm. We hypothesized that the N-terminus of BlaD encodes a signal sequence based on the signal sequence prediction within the first 18 amino acid residues (58, 59). We generated a truncated version of BlaD missing these first 18 residues (BlaDΔ18; p*blaD*Δ18). As shown in **Figure 5B**, the expression of BlaDΔ18 is unable to complement the absence of β-lactamase activity in the Δ*blaXD* mutant in a whole cell assay. qRT-PCR results shown in **Figure S9** confirm that *blaX* and/or *blaD* are expressed in the complemented strains, indicating that the absence of gene expression is not the cause of the lack of observable β-lactamase activity. This suggested that BlaDΔ18 is either not translated, is an unstable or inactive protein, or is active but trapped in the cytosol and unable to hydrolyze nitrocefin.

All of the characterized β-lactamases in Gram-positive bacteria are membrane-bound enzymes, although many of these proteins are cleaved, resulting in a smaller, soluble form that can be found in culture supernatants (29, 31, 34, 36). These findings are consistent with the lack of a periplasmic space for β-lactamases accumulation in Gram-positive bacteria. To determine if a soluble form of BlaD is secreted into the culture medium, we performed a nitrocefin hydrolysis assay using culture supernatants. As shown in **Figure 6A and 6C**, neither the supernatants of Δ*blaXD* cells harboring p*blaD* or p*blaX*-*blaD*, nor the wild-type strains 630Δ*erm* or M120, react with nitrocefin, indicating that BlaD is not secreted into the medium. To confirm that BlaD is a membrane-associated enzyme, we lysed the cells and performed a nitrocefin hydrolysis assay using lysates containing cell debris (denoted as ‘lysates’) or the cleared cell lysates (denoted as ‘lysate filtrate’). **Figures 6B and 6D** show that when comparing the level of activity in the lysate to the lysate filtrate in strains containing a full-length *blaD*, 74-80% of the total β-lactamase activity is found in the cell debris, indicating that BlaD is associated with the cell surface. Furthermore, BlaDΔ18 activity is not associated with the cell surface, as demonstrated by the similar levels of activity in the lysate and the lysate filtrate (**Figure 6B**). This result indicates that BlaDΔ18 is an active, soluble form of BlaD that is trapped in the cytosol, and strongly suggests that the first 18 residues at the N-terminus of BlaD encode a signal sequence. Together, these results support the presence of a signal sequence that helps bring the protein to the cell surface.

**Figure 6.**
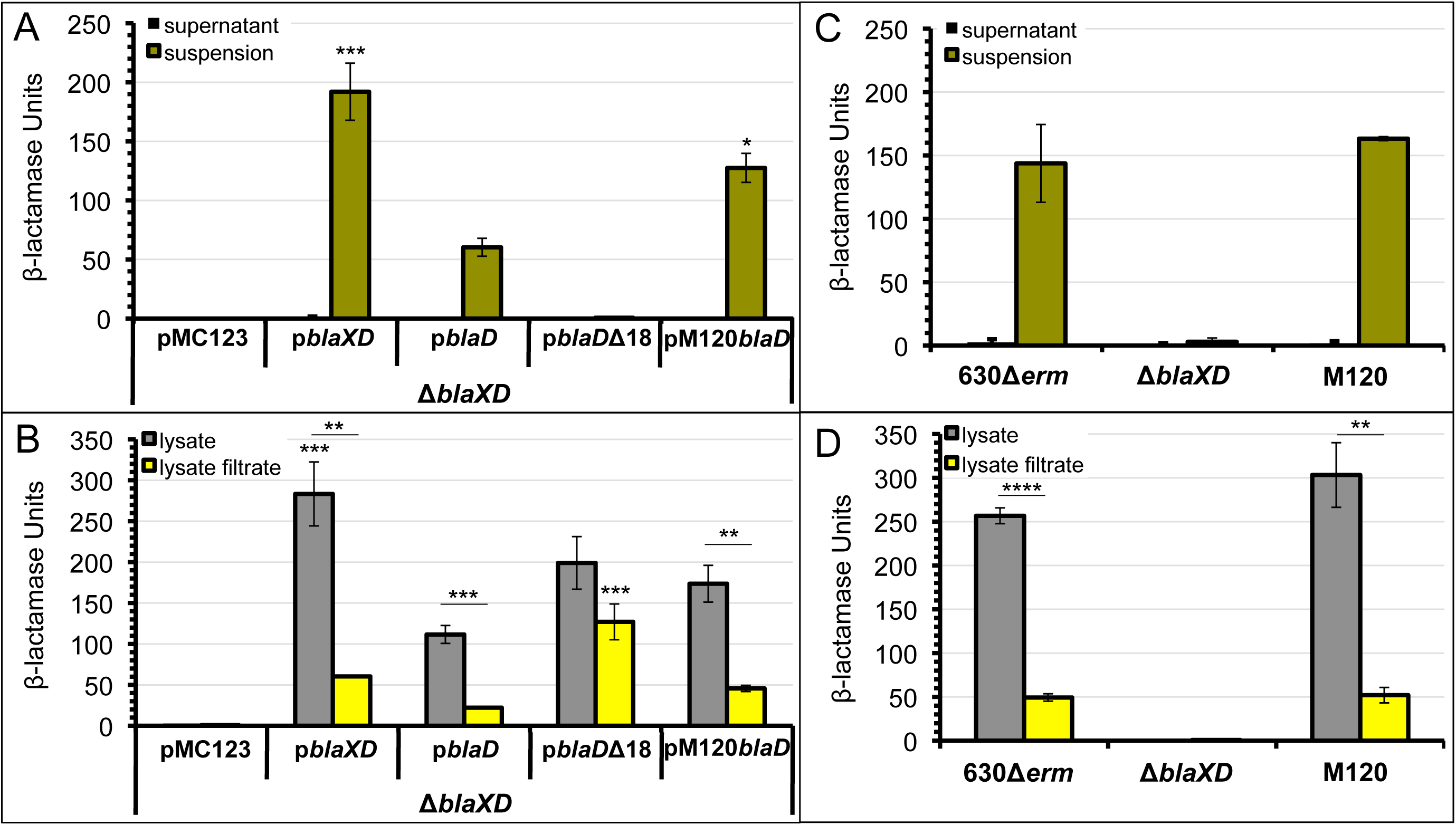
BlaD utilizes a signal sequence to act at the cell membrane. Δ*blaXD* (**A, B**) or 630Δ*erm*, Δ*blaXD*, and M120 (**C, D**) *C. difficile* were grown to mid-log phase in 2 µg/mL thiamphenicol and 2 µg/mL ampicillin and assayed for β-lactamase activity via a nitrocefin assay in **A, C**) supernatant or cell suspension and **B, D**) cell lysate or cell lysate filtrate. Δ*blaXD* pMC123 (MC 1400); Δ*blaXD* p*blaXD* (MC1399); Δ*blaXD* p*blaD* (MC1466); Δ*blaXD* p*blaD*Δ18 (MC1338); Δ*blaXD* pM120*blaD* (MC1494). Columns represent the means +/− SEM from at least three independent replicates. Data were analyzed by one-way ANOVA with Dunnett’s multiple comparisons test, compared to p*blaD* in A) and B) or 630Δ*erm* in C) and D), οr by a two-tailed unpaired student’s *t*-test, where indicated by bars. Absence of asterisk indicates no statistically significant difference found. Adjusted *P* values indicated by *≤0.05, ****<0.0001.

### BlaX aids in BlaD activity

Although BlaX is not necessary for BlaD activity (**Figure 5A, B**), *blaX* is conserved in many *C. difficile* strains. Thus, we examined whether BlaX enhances BlaD activity. The results shown in **Figure 6A and 6B** demonstrate that the presence of BlaX increases β-lactamase activity of the 630Δ*erm* BlaD two to three-fold, suggesting that BlaX plays a role in the function of BlaD. To investigate the activity of a BlaD from a *C. difficile* genome that lacks BlaX, we also complemented the Δ*blaXD* strain with *blaD* cloned from the M120 genome, under the M120 native promoter. **Figure 6A** shows that in cell suspensions of Δ*blaXD* complemented strains, the M120 BlaD (pM120*blaD*) exhibits two-fold higher activity than the 630Δ*erm* BlaD (p*blaD*). This result suggests that the M120 BlaD is superior to the 630Δ*erm* BlaD at translocating to the cell surface when BlaX is not present. However, M120 BlaD is only two-thirds as active as the 630Δ*erm* BlaXD complement (p*blaXD*). In lysed cells, the M120 BlaD β-lactamase activity levels are slightly higher than the 630Δ*erm* BlaD (**Figure 6B**). Interestingly, the wild-type strains 630Δ*erm* and M120 exhibit similar β-lactamase activity levels in both cell suspension and lysate samples, indicating that their overall efficacy is comparable (**Figure 6C and D**). Together, these results demonstrate that in 630Δ*erm*, BlaX enhances BlaD activity, while in M120, β-lactamase activity is not dependent on BlaX. Finally, because the M120 BlaD does not fully complement the Δ*blaXD* strain, the N-terminal sequence variability of the BlaD proteins likely plays a role in strain-dependent translocation of BlaD to the cell surface.

### The *bla* operon is regulated by BlaIR

Transcription of most β-lactamase genes in Gram-positive bacteria is regulated by the two-component BlaRI system (60-62). The *C. difficile* genome encodes several orthologs of the two genes that make up this system, *blaI* and *blaR.* In other bacteria, BlaR is a sensor that is activated upon β-lactam binding (63). Activated BlaR cleaves the BlaI repressor, which is bound as a dimer to the *bla* operon promoter in the absence of β-lactams (64-66). Once cleaved, BlaI can no longer bind to the *bla* promoter, thus allowing for active transcription. Two candidate orthologs *CD0471* (*blaI*) and CD0470 (*blaR*) are located 11 kb downstream of the *blaXD* operon. To determine if these *blaIR* orthologs regulate the *blaXD* operon in *C. difficile*, we created an insertional disruption in *blaI*. **Figure S10** shows that transcription of *blaI* and *blaR* are decreased in the *blaI*::*erm* mutant, confirming that *blaI* and *blaR* are organized in an operon, as is consistent with other bacteria. As shown in **Figure 7**, in the absence of β-lactams, *blaX* and *blaD* are transcribed at high levels in the *blaI*::*erm* mutant, as compared to the wild-type 630Δ*erm* strain. These results confirm that BlaI acts as a repressor of the *bla* operon. Further, the induction of *blaXD* in β-lactams in the wild-type strain, but not in the mutant, strongly suggests that BlaI repression is relieved by the presence of β-lactams in wild-type strain. To verify that relief of BlaI repression results in β-lactamase production, we performed a nitrocefin hydrolysis assay on the *blaI*::*erm* mutant. **Figure 5C** confirms that the absence of BlaI results in active β-lactamase, independent of β-lactam presence. Together, these results show that *C. difficile* encodes a BlaRI system that represses *bla* transcription in the absence of β-lactams. Efforts to complement *blaIR* resulted in poor growth of *E. coli* mating strains, as well as *C. difficile*, and were not successful.

**Figure 7.**
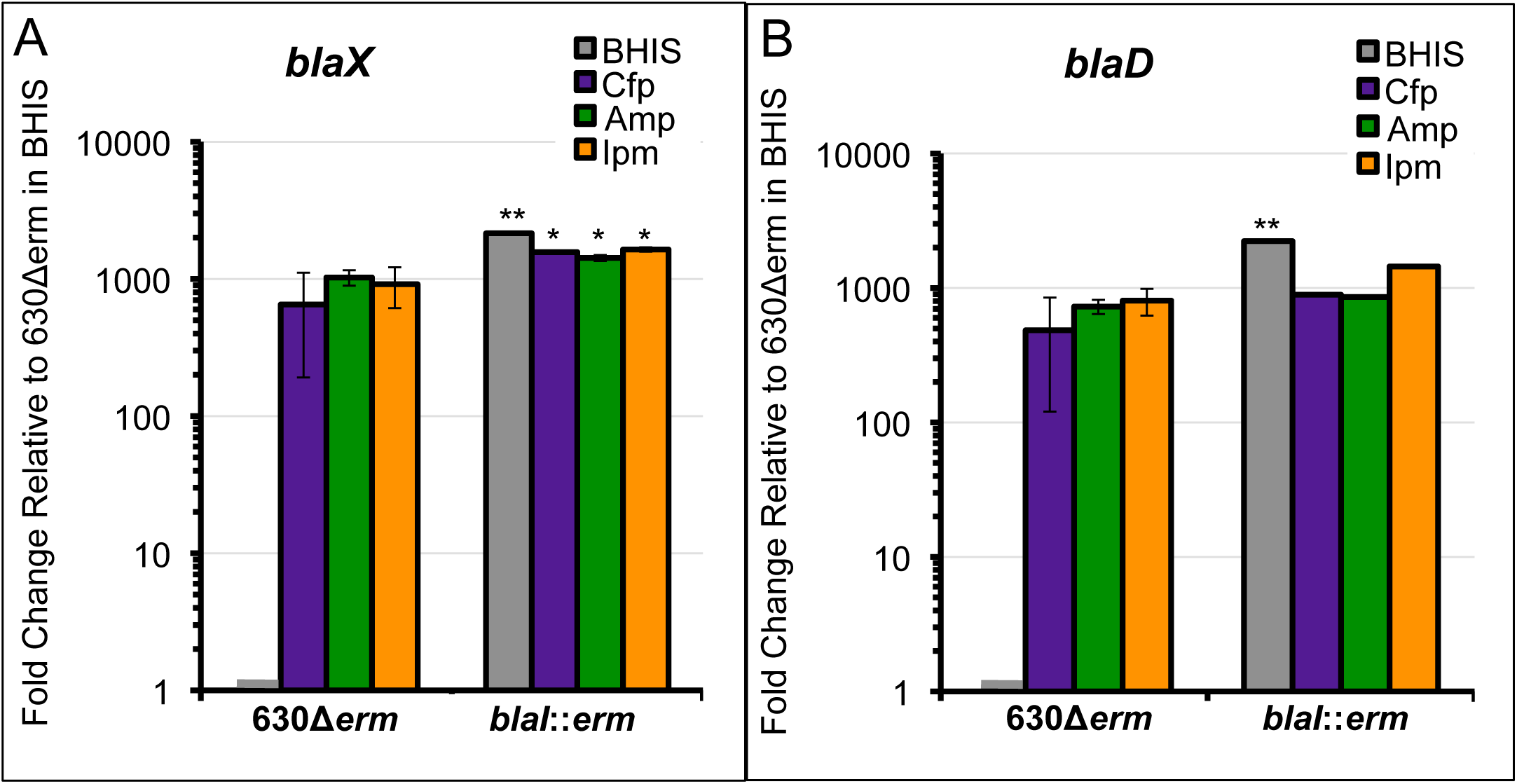
*blaXD* is derepressed in the *blaI*::*erm* strain. qRT-PCR was performed to measure expression of **A**) *blaX* and **B**) *blaD* in *C*. *difficile* 630Δ*erm* and *blaI*::*erm* strains grown to mid-log in BHIS media with or without β-lactam (Cfp: cefoperazone 60 µg/mL; Amp: ampicillin 2 µg/mL; Ipm: imipenem 1.5 µg/mL). mRNA levels are normalized to expression levels in 630Δ*erm* in BHIS alone. Columns represent the means +/− SEM from three independent replicates. Data were analyzed by one-way ANOVA with Dunnett’s multiple comparisons test, compared to expression in 630Δ*erm* without antibiotic. Adjusted *P* values indicated by *≤0.05, **≤0.005.

To further confirm that the BlaRI system regulates the *bla* operon and to define its contribution to ampicillin resistance, we examined the growth of the *blaI*::*erm* mutant in multiple β-lactams. **Figure 8A** illustrates that growth of the *blaI* mutant is not significantly different than the wild-type 630Δ*erm* strain in the presence of cefoperazone. However, growth of the *blaI* mutant is significantly improved in the presence of ampicillin, as compared to 630Δ*erm* (**Figure 8B**). Similarly, the *blaI*::*erm* mutant shows slightly impaired growth in imipenem, as compared to 630Δ*erm* (**Figure 8C**). These results show that BlaIR contributes to ampicillin and impenem resistance in *C. difficile* through regulation of the *bla* operon.

**Figure 8.**
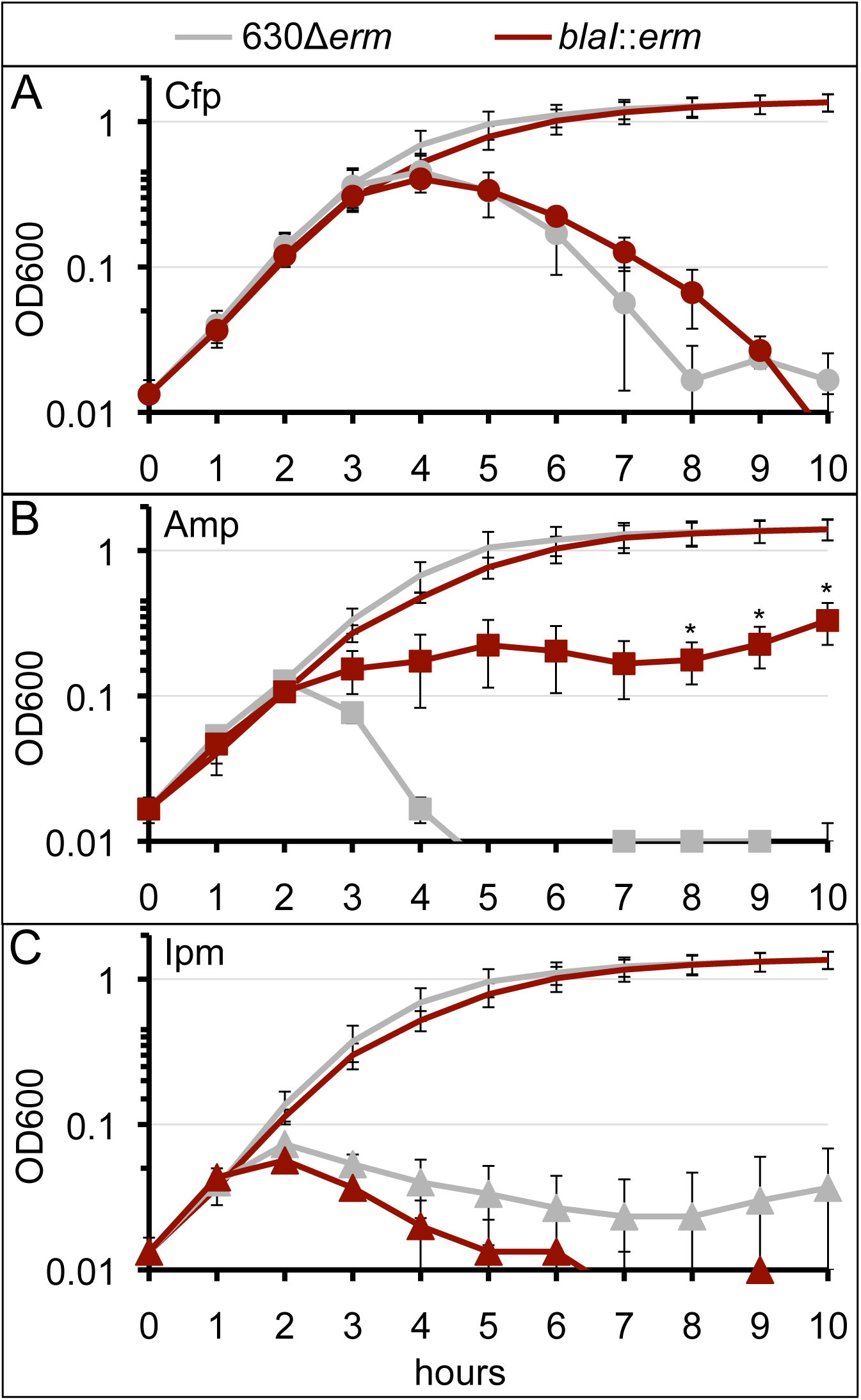
*blaI* grants resistance to ampicillin. *C. difficile* strains 630Δ*erm* (gray) and *blaI*::*erm* (MC985; red) were grown to mid-log, backdiluted to OD 0.05, and grown in BHIS (no marker) or BHIS supplemented (filled marker) with **A**) 60 µg/mL cefoperazone (Cfp), **B**) 4 µg/mL ampicillin (Amp), or **C**) 2 µg/mL imipenem (Ipm). Lines represent the means +/− SEM from three independent replicates. Data were analyzed by one-tailed paired Student’s *t*-test, compared to 630Δ*erm*. Adjusted *P* values indicated by *≤0.05.

## DISCUSSION

This study provides evidence for β-lactam-dependent expression of the β-lactamase, BlaD, in two strains of *C. difficile*, 630Δ*erm* and R20291, as well as activity of BlaD in both 630Δ*erm* and M120. The *blaD* gene is located in an operon with *blaX*, which encodes a putative membrane protein (**Figure S4**). Our data indicate that the promoter for the *blaXD* operon is located within a 300 nucleotide region located directly upstream of the *blaX* start codon (**Figure 3**). The high level of *blaD* and *blaX* expression in response to β-lactams far below MICs (**Figure S7**), indicate that the promoter of the *bla* operon is quite strong, in contrast to a previous report in which part of the *blaD* locus was expressed in a heterologous host (39).

Our work has demonstrated that BlaD is a β-lactamase that is only active under anaerobic (reducing) conditions (**Figure 1**). To our knowledge, no other anaerobic β-lactamases have been reported, which is not surprising given that β-lactamase assays are generally performed in the presence of oxygen (67, 68). This, however, may be one reason that so few β-lactamases have been identified in anaerobic, Gram-positive bacteria (69-72). Indeed, the addition of 0.2 mM DTT to the nitrocefin hydrolysis assays, or steady-state enzyme kinetics (39), allowed for observation of BlaD activity (**Figure 6**) by maintaining reducing conditions. Assaying β-lactamases from other anaerobic, Gram-positive bacteria under reducing conditions may lead to the identification of more anaerobic β-lactamases in other species.

Our data indicate that BlaD acts at the cell membrane, in accordance with other β-lactamases from Gram-positive bacteria (**Figure 6**). We have shown that BlaD likely contains a signal sequence at the N-terminus, which facilitates translocation of BlaD to the membrane. BlaD is not secreted into the environment, but remains associated with the cell surface (**Figure 6**). While the exact function of BlaX is unknown, the data demonstrate that BlaD activity is enhanced by the presence of BlaX (**Figure 6B**). BlaX has five predicted transmembrane domains, with an approximate 125 residue-long extracellular loop (73). Because the activity of BlaD is membrane-associated across all samples except BlaDΔ18, and BlaD activity in cell lysates lacking BlaX is 60% less than when BlaX is present, it is possible that BlaX interacts with BlaD in a way that makes BlaD more accessible to substrates on the cell surface. Nitrocefin hydrolysis assays showed that in cell lysates, the activity of full length BlaD (p*blaD*) is 45% less than BlaDΔ18 (**Figure 6B**). This suggests that either BlaD is cleaved at the N-terminus after translocation to the cell membrane, or BlaX helps to relieve a steric hindrance caused by insertion into the cell membrane. The absence of β-lactamase activity in cell supernatants does not support cleavage of BlaD, unless BlaD remains anchored to the cell membrane after cleavage, which is unlikely due to the absence of a canonical lipobox immediately downstream of the signal peptide (74).

To date, only one other published β-lactamase is reported to be co-transcribed with a membrane protein (75). This membrane-bound β-lactamase, PenA, found in the Gram-negative *Burkholderia psuedomallei*, is encoded in an operon with *nlpD1*, a gene annotated as an outer membrane lipoprotein and thought to be involved in cell wall hydrolytic amidase activation (76). However, *C. difficile* does not contain an outer membrane, and *nlpD1* is not homologous with *blaX*. Analysis of the *blaD* locus in the *C. difficile* strain M120, which does not contain a full *blaX* coding sequence, revealed regions of partial homology to the 5’ and 3’ ends of *blaX*, located between the promoter and the *blaD* start codon. This suggests that over the course of evolution of *C. difficile*, the majority of this gene was deleted. A search of the rest of the M120 genome revealed no other proteins similar to BlaX, further supporting the model that in many *C. difficile* strains, BlaX is not necessary for sufficient BlaD activity. However, the superior activity levels of M120 BlaD (**Figures 6A and 6B**), the 74% of cell surface-associated activity of M120 BlaD (**Figure 6B**), as well as the equal levels of β-lactamase activity of the 630Δ*erm* strain compared to M120 (**Figure 6D**), suggest that M120 likely has a different mechanism of translocation.

We have shown that the *bla* operon confers resistance to ampicillin and is regulated by the BlaRI system in *C. difficile* (**Figures 5, 8**). Disruption of *blaI* resulted in constitutive expression of *blaX* and *blaD* (**Figure 7**), which resulted in improved growth in ampicillin (**Figure 8**), supporting the model that BlaI is a direct repressor of the *bla* operon. We identified a 52-nucleotide region of dyad symmetry in the promoter of the *bla* operon, which contains a canonical BlaI binding site, supporting the model of BlaI-P*blaX* binding, but does not rule out other binding partners. Our results align with previously reported data that BlaD confers resistance to penicillins (39). The discrepancy of the MIC values versus the growth curves can be attributed to the exact nature of a growth curve. Further investigation is needed to fully define the mechanisms of β-lactam resistance in *C. difficile*. Identification and characterization of the additional β-lactam resistance mechanisms may aid in preventing *C. difficile* infections and recurrence in the future.

## ACKNOWLEDGEMENTS

We thank members of the McBride lab and the dissertation committee of B.K.S. for helpful suggestions and discussions throughout the course of this work. This research was supported by the U.S. National Institutes of Health (NIH) through research grants AI116933 and AI121684 to S.M.M., and training grant AI106699 to S.A. The content of this manuscript is solely the responsibility of the authors and does not necessarily reflect the official views of the National Institutes of Health.

## SUPPLEMENTAL FIGURE LEGENDS

**Figure S1. DNA cloning and vector details.**

**Figure S2. The putative β-lactamase gene, *CD0458 (CDR20291_0399)*, is induced by β-lactams.** Putative β-lactamase genes in strains **A**) 630Δ*erm* and **B**) R20291 were measured for relative expression to the housekeeping gene, *rpoC*, in β-lactams via qRT-PCR (Cfp: cefoperazone 50 μg/mL; Amp: ampicillin 2 μg/mL; Ipm: imipenem 1.5 μg/mL). mRNA levels are normalized to expression levels in BHIS alone. Columns represent the means +/− SEM from three independent replicates. Data were analyzed by one-way ANOVA with Dunnett’s multiple comparisons test, compared to no antibiotic. Adjusted *P* values indicated by *≤0.05, ***≤0.001.

**Figure S3. The putative β-lactamase, *CDR20291_0399*, and its upstream gene are induced by β-lactams. A**) The putative β-lactamase gene *CDR20291_0399* is 27 bp downstream of the putative membrane protein, *CDR20291_0398*. **B**) Relative expression of each gene was measured via qRT-PCR. *C. difficile* strain 630Δ*erm* was grown to mid-log in BHIS medium supplemented with sub-inhibitory concentrations of β-lactams (Cfp: cefoperazone 50 μg/mL; Amp: ampicillin 2 μg/mL; Ipm: imipenem 1.5 μg/mL). mRNA levels are normalized to expression levels in BHIS alone. Columns represent the means +/− SEM from three independent replicates. Data were analyzed by one-way ANOVA with Dunnett’s multiple comparisons test, compared to no antibiotic. Adjusted *P* values indicated by *≤0.05, **≤0.01.

**Figure S4. *blaX* and *blaD* form the *bla* operon. A**) PCR was performed using a forward primer (oMC1184) within *blaX* and the reverse primer (oMC1185) within *blaD* (*CD0457* and *CD0458* in 630Δ*erm* and *CDR20291_0398* and *CDR20291_0399* in R20291). **B**) cDNA was created from *C. difficile* strains 630Δ*erm* and R20291 treated with 2 μg/mL ampicillin. gDNA: genomic DNA from each strain served as a positive control; −RT: RNA from a reverse transcription reaction lacking enzyme served as a negative control.

**Figure S5. Analysis of gene expression of mutants *blaX*::*erm* and Δ*blaXD*.** Relative expression of **A**) *blaX* and **B**) *blaD* in 630Δ*erm* compared to *blaX*::*erm* (MC905), and Δ*blaXD* (MC1327) was measured via qRT-PCR. *C. difficile* was grown to mid-log in BHIS media supplemented with sub-inhibitory concentrations of β-lactams (Cfp: cefoperazone 60 μg/mL, Amp: ampicillin 2 μg/mL, and Ipm: imipenem 1.5 μg/mL). mRNA levels are normalized to expression levels in 630Δ*erm* in BHIS alone. Columns represent the means +/− SEM from three independent replicates. Data were analyzed by one-way ANOVA with Dunnett’s multiple comparisons test, compared to expression in 630Δ*erm* without antibiotic. Adjusted *P* values indicated by *≤0.05, ****<0.0001.

**Figure S6. *blaXD* transcription is modestly induced by vancomycin and polymyxin B.** Relative expression of each gene was measured via qRT-PCR. *C. difficile* strain 630Δ*erm* was grown to mid-log in BHIS medium supplemented with sub-inhibitory concentrations of cell wall targeting antimicrobials (Van: vancomycin 0.75 μg/mL, PmB: polymyxin B 75 μg/mL, Lys: lysozyme 1 mg/mL, Nis: nisin 7.5 μg/mL, LL-37 2 μg/mL, and Kan: kanamycin 250 μg/mL). mRNA levels are normalized to expression levels in BHIS alone. Columns represent the means +/− SEM from four independent replicates. Data were analyzed by one-way ANOVA with Dunnett’s multiple comparisons test, compared to expression in 630Δ*erm* without antibiotic. No statistically significant values found.

**Figure S7. The *bla* operon exhibits dose dependent expression for different classes of β-lactams.** Relative expression of *blaX* and *blaD* in 630Δ*erm* was measured using qRT-PCR. *C. difficile* was grown to mid-log in BHIS medium supplemented with increasing sub-inhibitory concentrations of **A**) cefoperazone (μg/mL: 3.125, 6.25, 12.5, 25, 50), **B**) ampicillin (μg/mL: 0.25, 0.5, 1, 1.5, 2), or **C**) imipenem (μg/mL: 0.125, 0.25, 0.5, 1, 1.5). mRNA levels are normalized to expression levels in 630Δ*erm* in BHIS alone. Columns represent the means +/− SEM from three independent replicates. Data were analyzed by one-way ANOVA with Dunnett’s multiple comparisons test, comparing to expression with lowest concentration antibiotic. Adjusted *P* values indicated by *≤0.05, ****<0.0001.

**Figure S8*. C. difficile* strain M120 displays β-lactamase activity. A**) Alignment of BlaD proteins from strains 630Δ*erm* and M120 via SerialCloner. The blue lines indicate the signal peptides predicted by Signal-3L 2.0 (58). The black triangle represents the site of truncation in the p*blaD*Δ18 (pMC811) construct. The red boxes indicate transmembrane domains predicted by Phobius (73). **B**) Nitrocefin hydrolysis assay of *C. difficile* strains 630Δ*erm* and M120. Strains were grown to mid-log in BHIS media only (a, c) or with 2 μg/mL ampicillin (b, d). Color change from yellow to red indicates cleavage of nitrocefin.

**Figure S9. Expression of *blaX* or *blaD* from Δ*blaXD* complemented strains.** qRT-PCR was performed to examine expression of **A**) *blaX* or **B**) *blaD* from a plasmid maintained in Δ*blaXD (*MC1327) grown to mid-log in BHIS media supplemented with 2 ug/mL ampicillin. mRNA levels are normalized to expression levels in Δ*blaXD (*MC1327) expressing an empty vector (pMC123) in BHIS alone. p*blaXD*: pMC867; p*blaD*: pMC897; p*blaD*Δ18: pMC811. Columns represent the means +/− SEM from three independent replicates. Data were analyzed by one-way ANOVA with Dunnett’s multiple comparisons test, comparing to expression without antibiotic. Absence of asterisk indicates no statistically significant difference found. Adjusted *P* values indicated by *≤0.05, **<0.0001.

**Figure S10. *blaIR* is derepressed but disrupted in the *blaI*::*erm* strain. A**) blaI was disrupted by an insertion. qRT-PCR was performed to measure expression of **B**) *blaI* and **C**) *blaR* in *C*. *difficile* 630Δ*erm* and *blaI*::*erm* strains grown to mid-log in BHIS media with or without β-lactam (Cfp: cefoperazone 60 μg/mL; Amp: ampicillin 2 μg/mL; Ipm: imipenem 1.5 μg/mL). mRNA levels are normalized to expression levels in 630Δ*erm* in BHIS alone. Columns represent the means +/− SEM from three independent replicates. Data were analyzed by one-way ANOVA with Dunnett’s multiple comparisons test, compared to expression in 630Δ*erm* without antibiotic. Adjusted *P* values indicated by *≤0.05, ****<0.0001.

## SUPPLEMENTAL TABLE LEGEND

**Table S1. MIC values for 630Δ*erm*, *blaX*::*erm*, and Δ*blaXD* strains.** MIC values were determined for strains 630Δ*erm*, *blaX::erm* (MC905), and Δ*blaXD* (MC1327) in Cfp (cefoperazone), Amp (ampicilin), and Ipm (imipenem) using liquid broth dilution. Values represent the highest MIC value of three biological replicates.

